# Protein folding stability estimation with explicit consideration of unfolded states

**DOI:** 10.1101/2025.02.10.637420

**Authors:** Heechan Lee, Yugyeong Cho, Jeongwon Yun, Martin Steinegger, Ho Min Kim, Hahnbeom Park

**Author notes:** Correspondence: Hahnbeom Park, Ho Min Kim.

## Abstract

Folding stability is a critical requirement for the vast majority of proteins. Computational methods suggested to date for the absolute folding stability (Δ*G*) prediction – including those driven from protein structure prediction AIs – show clear limitations on reproducing quantitative experimental values. Here we present IFUM, a deep neural network that jointly estimates Δ*G* and the equilibrium ensemble of folded and unfolded states represented by their residue-pair distance probability distributions. This joint learning considerably enhances the Δ*G* prediction accuracy against the scenario where Δ*G* prediction was learned alone. To improve the model, we extend the dataset beyond previous related works to include the Mega-scale small proteins and disordered proteins for training as well as wild-type natural proteins with sizes up to 869 residues for validation. We show that IFUM is robust to various protein types and sizes, and is capable of accurately predicting more general types of mutational effects such as sequence insertions or deletions. The applicability of IFUM is demonstrated through two real-world design challenges. First, for blind-tested protein engineering scenarios containing many sequence substitutions and insertions, good correlation with experimental melting temperatures (T_m_) is observed. Second, for the *de novo* design selection, IFUM shows considerably improved performance over broadly used AlphaFold-based metrics. IFUM is a free software available at github.com/HParklab/IFUM and also through Google Colab.

## Introduction

A majority of proteins function by folding into unique three-dimensional structures^1^. Because naturally evolved proteins are often only marginally stable in their unfolding free energy (Δ*G*_unfolding_, or Δ*G* hereafter)^2,3^, a small shift in the folding stability can result in a crucial impact. For instance, structure-disrupting mutations with poorer Δ*G* can lead to protein misfolding and aggregation which can eventually cause human diseases^4,5^ or poor protein expressions^6^. Thus, accurate Δ*G* quantification is highly demanded in many biotechnological applications, such as therapeutic development or biocatalysis^7,8^.

Measuring the Δ*G* of a protein experimentally presents significant challenges due to several factors. These measurements often require extrapolation from non-native conditions, such as high concentrations of denaturants or elevated temperatures, which can introduce inaccuracies^9^. Furthermore, the process is typically labor-intensive, necessitating the purification of each protein variant, even with the aid of automated methods. The resulting experimental data are also frequently inhomogeneous, stemming from variations in methods and experimental conditions across different studies, making direct comparisons and comprehensive analyses difficult^9^.

Computational estimation of Δ*G*, which can significantly reduce such experimental cost, has been studied from various perspectives. One of the most popular methods these days is to approximate it using the confidence estimate from structure prediction results. For instance, AlphaFold^10,11^ or ESMFold^12^ provide metrics called plDDT and pTM, which are confidence estimates of structural accuracy of predicted structures learned by separate modules. Although not explicitly learned for protein stability, it has been shown that those metrics can be effective for filtering out unstable designed proteins^13^, and then became a popular computational filter in various design studies^13,14^. However, these methods are not designed to estimate Δ*G* and therefore show limitations in providing a quantitative estimate. Similarly, the pseudolikelihood assigned to a sequence by protein language models (PLMs) or inverse folding models has been explored as a proxy for protein stability^15^. In parallel, considerable efforts have been made to build networks for the estimation of ΔΔ*G,* the change in stability upon mutation (Δ*G*_mutant_ – Δ*G*_wildtype_)^16,17^. However, despite their utility for protein engineering, these methods provide little insight to the absolute protein stability, and are generally limited to single point mutations.

We reasoned that the challenge of Δ*G* prediction originates from the ambiguity of the sequence-to-structure relationship at unfolded states. Because Δ*G* is by definition the gap between *G*_folded_ and *G*_unfolded_, to accurately quantify Δ*G*, it is very natural to consider the unfolded state along with the folded state. In fact, sequences not only influence the thermodynamic characteristics of folded states but also that of unfolded states; for instance, unfolded states of poly-Leu and poly-Lys peptides cannot be equal to each other. This sequence-dependence to unfolded states was either modeled using a linear model called “reference state energy”^18^, which was quite effective yet too simple to capture the complex nature of unfolded states, or was unmodeled and left as a “null state” regardless of protein sequences in many deep learning approaches^15,19^. We hypothesized that the key question in this endeavor should be how to explicitly abstract the unfolded state into a deep learning architecture to effectively represent the protein unfolded state.

In this work, we introduce IFUM (*In silico* evaluation of unfolding Free energy with Unfolded state ensemble Modeling, pronounced [ip͈ɯm]), a deep learning model that addresses the challenge of accurate Δ*G* prediction by explicitly abstracting the unfolded state ensemble. Conceptually speaking, IFUM makes two hypotheses to abstract the unfolded state representation: first, proteins fold via a two-state folding model^20^, and second, that the myriad conformations within the unfolded state ensemble can be effectively simplified and represented as an ensemble-averaged distance map using the Flory random coil model^21^. The Flory random coil model, a fundamental concept in polymer physics, describes an ideal polymer chain whose conformation is statistically averaged as a random walk, thus providing a simplified yet powerful framework for modeling the highly diverse and disordered ensemble of protein unfolded states^22^. By employing these principles, IFUM estimates Δ*G* and the relative thermodynamic preferences of a given sequence at the average structures of folded and unfolded states represented as residue-pairwise distance maps, which corresponds to the “unfolding free energy” definition: Δ*G* = –R*T* ln([*U*]/[*F*])^20^. IFUM demonstrates that this approximation works for accurately predicting Δ*G* for various types of proteins, and also for estimating ΔΔ*G* from sequence deletions or insertions that previous ΔΔ*G*-centric methods were not capable of.

## Results

### Overview of IFUM

As highlighted above, the key concept of IFUM is to abstract the conformations of both folded and unfolded states of proteins. We use the residue-pair Cα distance distribution histogram (distogram) for this purpose. IFUM takes two types of inputs from pre-trained models: sequence-based and structure-based ones. The sequence embedding is obtained from ProtT5^23^. The folded state structural embedding is generated using ESM-IF1^24^ on the structure predicted by deep-learning based structure prediction tools such as ESMFold^12^. This structural embedding and the mean pairwise Cα distances (as a form of one-hot encoded distogram with 21 bins with a range of 2 to 42 Å) of the folded state structure are fed as structural inputs (**Fig. 1A, Fig. S1**). No specific input was provided for the unfolded state because the Flory model only depends on the residue-pair sequence separation.

**Fig. 1:**
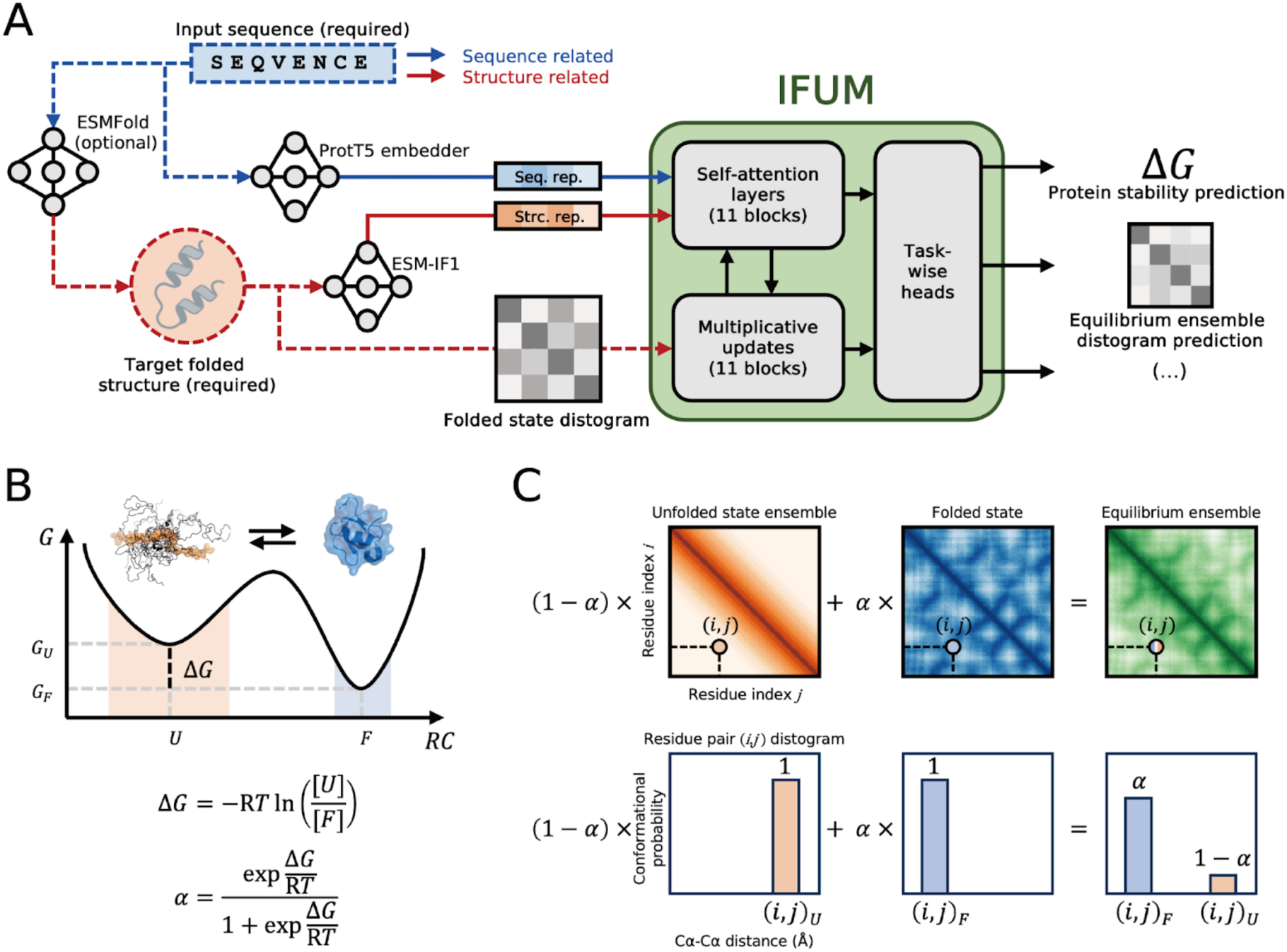
Overview of IFUM framework and key concepts. (A) Schematic illustration of the IFUM model. Sequence embeddings from ProtT5 (blue solid arrow; seq. rep.), structure embeddings from ESM-IF1 (red solid arrow; str. rep.), and a residue-pair Cα distogram (distance histogram) from ESMFold (red dashed arrow; folded state distogram) are fed into a main module with task-specific heads. (B) Equilibrium ensemble derivation. Energy landscape diagram illustrating the unfolded (*U*, *G_U_*) and folded (*F*, *G_F_*) states separated by Δ*G*, with protein schematics and formulas for Δ*G* and the folded state population α. (C) Derivation of the equilibrium ensemble distogram. Top: Distograms for the unfolded and folded states are weighted by (1−α) and α respectively, and then summed. Example residue-pair (*i*,*j*) are highlighted. Bottom: Cα-Cα distogram for a residue pair (*i,j*). Bar charts show unfolded (left) and folded (middle) distograms, weighted by conformational probabilities (1−α) and α, to give the equilibrium ensemble (right) distogram (see Methods).

Using the sequence and the folded-state structural information, a transformer-based module jointly updates these embeddings and the distogram inspired by AlphaFold2’s Evoformer^10^. The network jointly estimates target objectives by applying separate heads: Δ*G*, ensemble distogram, and an auxiliary head for sequence recovery. The Δ*G* head predicts per-residue Δ*G*, a vector with the same length as a protein, which is summed up to predict the net Δ*G* of a protein. Protein Δ*G* labels are derived from experimental measurements (Mega-scale^20^) or set as “less than 0.5 kcal/mol” for disordered proteins (DisProt^25^). The distogram head predicts the probability distribution of distances for all residue pairs given the sequences and the estimated folded structure distances. For the ensemble distogram labels, mean residue distances at the folded state derived from the predicted structure and the unfolded state derived from the Flory model are one-hot encoded into a distogram form, with the ratio derived from the Boltzmann distribution (**Fig. 1B, C**). By doing so, the network can effectively learn to what extent the given sequence can accommodate a denatured structure with respect to the folded structure.

IFUM was trained on a dataset composed of two distinct sources: a subset of 648,650 proteins (30-80 amino acids in length) from the Mega-scale dataset^20^, which features Δ*G* values inferred from cDNA display proteolysis, and a set of 3,219 disordered proteins from the DisProt database^25^ with at most 70 amino acids. For a rigorous test-only evaluation, we compiled several independent datasets, including i) 6,007 sequences from the CATH database^26^ with lengths up to 869 amino acids, ii) 26 protein-engineered sequences with experimentally measured melting temperatures (T_m_) (12 IFN-λ^14^, 8 IL-10, and 6 UGT76G1^8^), iii) a manually curated literature set of 40 unique wild-type sequences with experimental Δ*G* values from the S669 dataset^24^, and iv) a manually curated literature set of 413 *de novo* designed sequences from five distinctive folds with their corresponding experimental expression data^27–33^ (**Table S1**). To ensure no data leakage, none of the sequences in these test sets shared more than the sequence identity of 0.30 with any data in the Mega-scale training set. Further details on the model architecture, inputs, Δ*G* labels, unfolded state ensemble modeling, and dataset curations are available in the **Methods**.

### IFUM can predict unfolding free energy within a low mean error

We first tested the accuracy of IFUM predictions on the Mega-scale test set. Testing on this set offers an objective estimation of the network’s precision limit, at least for small proteins, at ideal conditions free from experimental disparities (e.g. pH, temperature, solvent, etc). IFUM achieved an RMSE (root mean squared error) of 1.16 kcal/mol and a PCC (Pearson correlation coefficient) of 0.78 when both labels and predictions are clamped to the dataset’s experimental dynamic range, [–1, 5] kcal/mol^20^ (**Fig. 2A**, more detailed statistical analyses provided in **Table S2**).

**Fig. 2.**
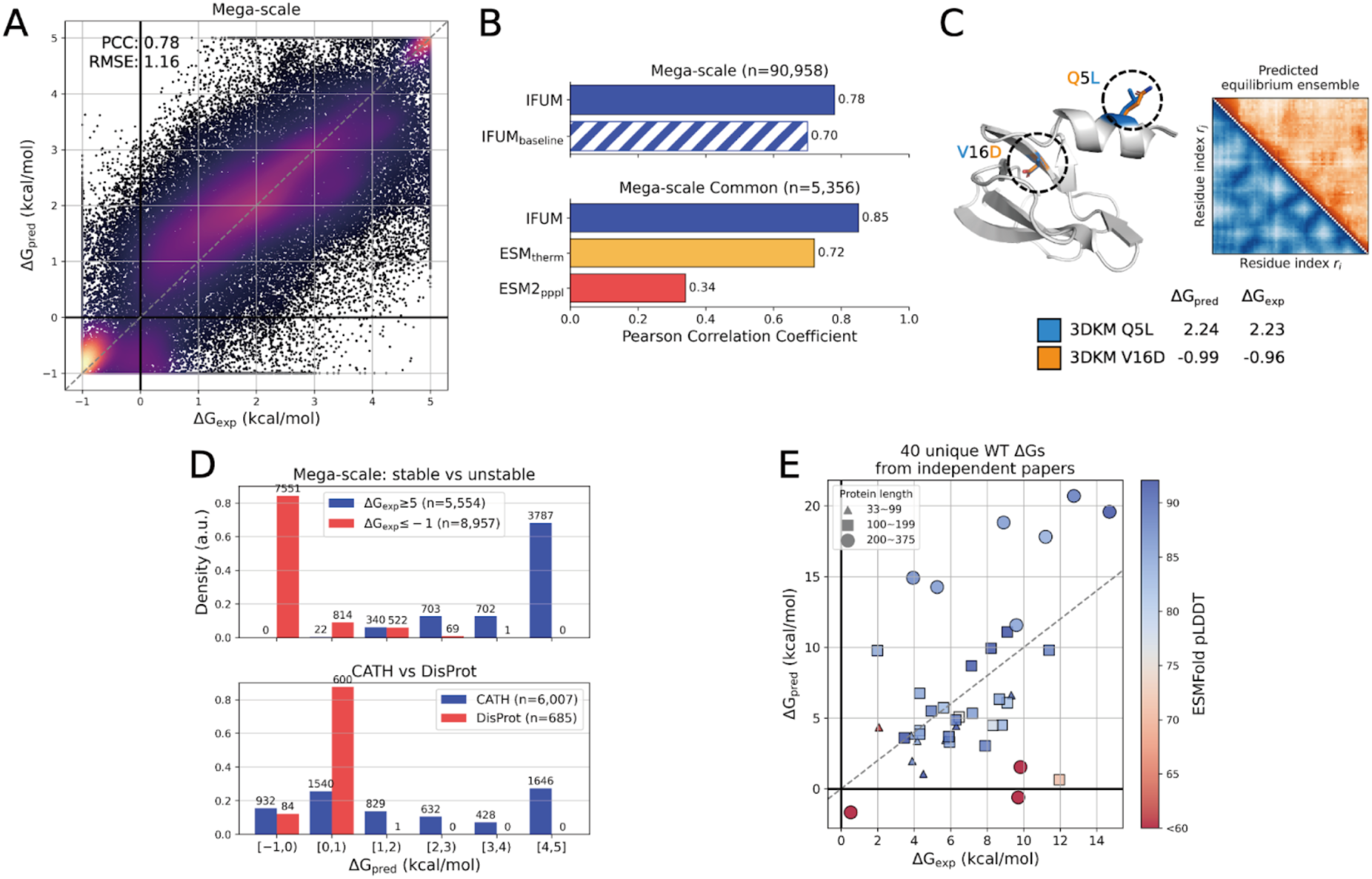
IFUM performance on Mega-scale and other datasets. (A) Scatter plot of IFUM-predicted Δ*G* (Δ*G*_pred_) versus experimental Δ*G* (Δ*G*_exp_) for the Mega-scale test set. Δ*G* values are clamped between –1 and 5 kcal/mol, according to the experimental dynamic range reported ^20^. Color intensity indicates point density (bright: high density; dark: low density). (B) Pearson correlation coefficient comparisons. Top: IFUM compared to an IFUM baseline model trained without unfolded state modeling on the Mega-scale test set. Bottom: IFUM compared to ESM_therm_ and ESM2 pseudo-perplexity (ESM2_pppl_) on the Mega-scale Common (see Methods). (C) Example predictions for HECTD1 CPH domain (PDB: 3DKM) mutants Q5L and V16D. Left: Overlay of ESMFold-predicted structures for Q5L (blue) and V16D (orange) mutants, showing their Δ*G*_pred_ and Δ*G*_exp_ values. Mutation sites (Q5L and V16D) are indicated with a dotted circle. Right: Corresponding predicted equilibrium ensemble distograms (orange: V16D, blue: Q5L). (D) Histograms comparing Δ*G*_pred_ values on different subsets. Top: Stable (Δ*G*_exp_ ≥ 5 kcal/mol) and unstable (Δ*G*_exp_ ≤ –1 kcal/mol) subsets of the Mega-scale test set. Bottom: A comparison between CATH and DisProt datasets. The x-axis labels denote specific ranges of Δ*G*_pred_ values, where parentheses indicate an exclusive boundary and square brackets indicate an inclusive boundary. (E) A scatter plot of Δ*G*_pred_ versus Δ*G*_exp_ for 40 unique wild-type proteins from literature data. The marker size corresponds to protein length, and the marker color corresponds to the ESMFold plDDT score. The dashed line indicates a perfect correlation.

To see how this compares to other existing tools, we ran ESM_therm_^19^, a fine-tuned version of ESM2 for Δ*G* prediction, and collected the ESM2 sequence pseudolikelihood (see **Methods**)^12^. For a fair comparison, we used the sequences that overlap between two test sets from IFUM and ESM_therm,_. This resulted in a common test set (noted as Mega-scale Common hereafter) of 5,356 sequences from the original Mega-scale dataset. With this particular smaller test set (Mega-scale Common), IFUM achieved a PCC of 0.85, outperforming both ESM_therm_ and ESM2 (0.72 and 0.34, respectively) (**Fig. 2B***, bottom*).

### Training IFUM to predict unfolded state ensemble improves Δ*G* prediction

Then, we confirmed whether this improvement resulted from the incorporation of unfolded state ensemble modeling. Training a simpler version of IFUM that does not contain any unfolded-state-related aspects (IFUM_baseline_), the PCC dropped from 0.78 to 0.70 (**Fig. 2B***, top*), with RMSE change from 1.16 to 1.39 kcal/mol when clamped (**Table S2**). Inspecting the equilibrium ensemble distogram prediction, it was highly consistent with the Δ*G* prediction; while stable proteins had ordered distograms resembling those from folded states, unstable ones had fuzzy distograms close to mixtures of folded and unfolded distograms (**Fig. 2C**). More examples showing the consistency between the model’s predicted equilibrium ensemble and Δ*G* are provided in **Fig. S2**.

We further investigated ablation studies to reveal the importance of key components of the network (**Table S2**). Removing the triangle multiplicative updates^10^ or replacing all differential self-attention layers^34^ with conventional self-attention significantly reduced performance. Critically, training IFUM without the equilibrium ensemble prediction objective resulted in a comparable performance decrease. This shows that, for the Δ*G* estimation, equilibrium ensemble modeling is as crucial as other well-established innovations in deep learning architectures.

### IFUM is broadly applicable to various types of proteins

Given the promising prediction accuracy on small proteins at highly controlled conditions (PBS, pH 7.4, T=298K), we next evaluated IFUM’s performance on more realistic problems. First, we asked about its discriminative power in distinguishing unstable (Δ*G*_exp_ ≤ –1 kcal/mol) versus stable (Δ*G*_exp_ ≥ 5 kcal/mol) sequences within the well-controlled Mega-scale test set (**Fig. 2D***, top*). The network was able to discriminate between those, giving a Welch’s t-test p-value << 0.001. Next, we moved on to a more challenging but realistic problem, which was to apply the same discrimination test on semi-labeled wild-type natural proteins. This dataset includes 6,007 folded sequences from CATH which contains structured domains from wild-type proteins, and 685 disordered proteins from DisProt (**Fig. 2D***, bottom*; see **Methods**). We found that the distributions of Δ*G*_pred_ values reflected the differences in the characteristics in two distinct datasets, giving a Welch’s t-test p-value << 0.001.

When investigating CATH domains with low Δ*G*_pred_ (i.e., unstable, < 1 kcal/mol), we observed a higher prevalence of solvent-exposed hydrophobic residues. Solvent-exposed hydrophobics can be commonly found in CATH domains that are located 1) at obligate oligomers or domain interfaces or 2) inside the membrane. To quantify this, we calculated the spatial aggregation propensity (SAP)^35^ using Rosetta (SapScoreMetric), where a higher SAP value indicates a greater propensity for aggregation. While average per-residue SAP of 0.4 to 0.5 is typically used to filter soluble proteins^36^, these “self-unstable-in-water” domains showed significantly higher per-residue SAP values (mean 1.08) (**Table S3**, **Fig. S3**). This exposure stems from artifacts in some CATH domains lacking their complete context (**Fig. S4**, **Fig. S5**). This analysis underscores the need for cautious IFUM application depending on the expected protein context.

We then tested its performance on labeled proteins of broader types, with sizes ranging from 33 to 375 residues. A set of 40 wild-type protein sequences was collected with their experimentally determined Δ*G* values^24^ (**Fig. 2E, Table S1**; see **Methods**). Across all 40 sequences, IFUM achieved a PCC of 0.47, which is poorer than that on the well-controlled data set. We found that the quality of folded state structure affects the prediction accuracy; restricting the analysis to the 34 sequences with plDDT_ESMfold_ > 80 improved the PCC to 0.61; with plDDT_ESMfold_ > 90 (16 sequences), PCC further increased to 0.73. These findings suggest that IFUM’s Δ*G* prediction accuracy is correlated with the confidence and quality of the input target folded state, with higher accuracy observed for proteins with well-defined structures. Further analysis between Δ*G* prediction and folded state quality is in the **Discussion**.

### IFUM can accurately predict ΔΔ*G* for various types of mutants

IFUM exhibited strong predictive performance for ΔΔ*G*, encompassing point/double and insertion/deletion (indel) mutants within the Mega-scale test set (**Fig. 3A**). Within the full Mega-scale test set, IFUM achieved a PCC of 0.81, 0.80, and 0.63 for point mutants, indels, and double mutants, respectively. To further benchmark this capability, we used the Mega-scale Common test set and compared IFUM against ThermoMPNN^16^ (ThermoMPNN-D^37^ for double mutants), ESM_therm_, ESM2, FoldX^38^ and Rosetta FastRelax and Cartesian ddg protocols^39,40^ (**Fig. 3C**). For Mega-scale Common indels, a comparison with ThermoMPNN was not applicable, IFUM demonstrated a clear advantage with a PCC of 0.76 over ESM_therm_, ESM2, FoldX, and Rosetta (0.66, 0.35, 0.22, and 0,01, respectively). For Mega-scale Common double mutants, IFUM achieved a PCC of 0.61, and the others 0.38, 0.70, 0.50, 0.50, and 0.78 (ThermoMPNN-D, ESM_therm_, ESM2, FoldX, and Rosetta, respectively).

**Fig. 3:**
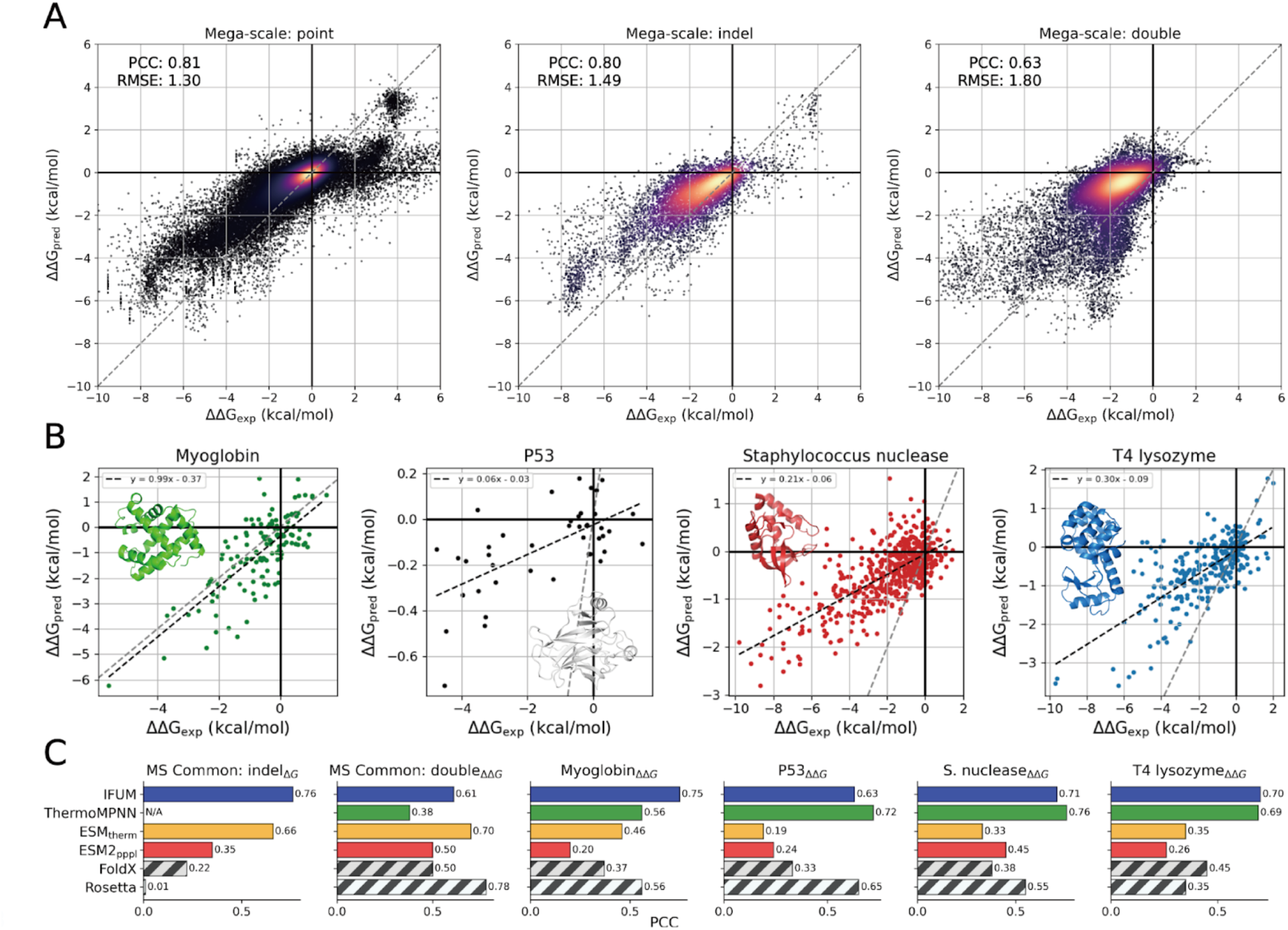
IFUM performance on various mutation types and comparison with other methods. (A) Scatter plots of IFUM-predicted ΔΔ*G* (ΔΔ*G*_pred_) versus experimental ΔΔ*G* (ΔΔ*G*_exp_) for different mutation types within the Mega-scale test set. Left: Point mutants. Middle: Insertion/deletion (indel) mutants. Right: Double mutants. Color intensity represents point density (bright = high density). (B) Scatter plots of ΔΔ*G*_pred_ versus ΔΔ*G*_exp_ for four case study proteins: Myoglobin, P53, S. nuclease, and T4 lysozyme. The dashed gray lines indicate perfect correlation, and the dashed black lines are linear regression lines. (C) Bar chart comparing Pearson correlation coefficients (PCC) for IFUM against ThermoMPNN, ESM_therm_, ESM2_pppl_, FoldX, and Rosetta. Comparisons are shown for the Mega-scale Common (MS Common) indel subset (Δ*G*), MS Common double mutant subset (ΔΔ*G*), and the four case study datasets (ΔΔ*G*).

Next, we evaluated the models on four case studies selected for single point mutation ΔΔ*G*: Myoglobin^41^, P53^42^, S. nuclease^43^, and T4 lysozyme^43^. IFUM achieved PCC values of 0.75, 0.63, 0.71, and 0.70, respectively (**Fig. 3B**), comparable to the ThermoMPNN specifically trained for single point mutations (0.56, 0.72, 0.76, and 0.69, respectively). Other methods (ESM_therm_ and ESM2) showed relatively poor performance in reproducing point mutation effects. Specific PCC values are reported in **Fig. 3C**, more details on double mutants ΔΔ*G* predictions are shown in **Fig. S6** and **Table S4**. Overall, IFUM was the only method among tests that robustly worked across various types of mutations ranging from single point mutations to indels.

In the following paragraphs, we demonstrate the practical utility of IFUM on two important protein design or engineering problems.

### Practical application 1. Protein stabilization engineering

It is quite common practice in protein engineering to introduce sequence length modifications or multiple-sequence substitutions. For instance, loops are often truncated for the protein stabilization engineering^8^ but are elongated instead when new functionality needs to be introduced^44^. Although structure prediction confidence estimations are broadly used as guidelines (e.g. filtering) for these processes^13^, their quantitative contribution remains unclear. Here, we demonstrate IFUM can be a good alternative to these structure prediction metrics.

We tested it on three protein engineering scenarios in which multiple sequence substitutions and/or sequence length modifications were introduced (**Fig 4**). For two targets (IFN-λ3 and IL-10), IFUM was blind tested in parallel with experimental human-driven engineering; for UGT76G1, IFUM was compared against previously reported data (per-target details in following paragraphs). Computational metrics were compared to experimental melting temperature (T_m_) values, which is known to well correlate with Δ*G* within the same protein domain^45,46^ (**Fig. S7**). Rosetta FastRelaxed AF3 models were used for the folded state structural input to the network.

**Fig. 4:**
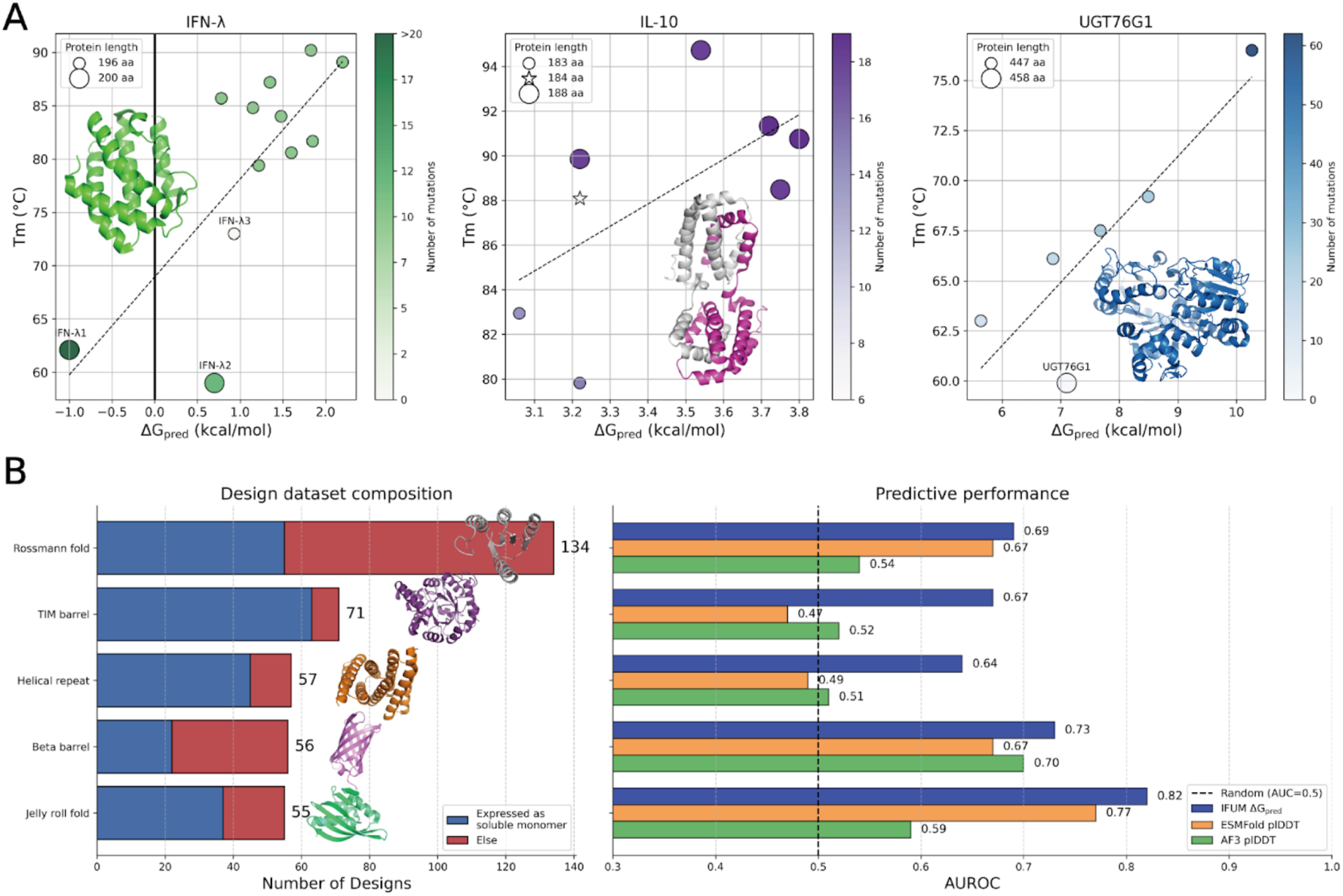
IFUM performance on real-world protein engineering and *in silico* screening applications. (A) Scatter plots of experimental melting temperature (T_m_) versus IFUM-predicted Δ*G* (Δ*G*_pred_) for IFN-λ, IL-10, and UGT76G1 sequence and backbone redesigns. The marker size corresponds to protein length, and the marker color corresponds to the number of mutations relative to each wild-type protein. Markers with the labels indicate the wild-type proteins. Rosetta FastRelaxed AF3 model structures were used in these scatter plots. The dashed line is a linear regression fit. (B) Performance on the *in silico* screening of designed proteins. Left: Composition of design datasets for five different protein folds, showing the number of computationally designed proteins that were experimentally found to express in E. coli in a soluble monomeric form (blue) or not (red). Right: Predictive performance, measured by the Area Under the Receiver Operating Characteristic curve (AUROC), for classifying non-expressing designs from expressing ones. The performance of IFUM’s predicted Δ*G*_pred_ (blue) is compared with the plDDT from ESMFold (orange) and AF3 (green).

The first group is engineered IFN-λ3 proteins to enhance its protein fitness and stability. In particular, the flexible loop harboring the thrombin cleavage site was excised and replaced with a structurally inpainted α-helix, which effectively shielded the adjacent hydrophobic patch and conferred resistance to proteolytic degradation. Thermal shift assays were performed to determine the T_m_ of 9 engineered IFN-λ3 variants and, for comparison, 3 wild-type IFN-λ isoforms (IFN-λ1, IFN-λ2, and IFN-λ3)^14^. The wild-type IFN-λ1 and IFN-λ2 consists of 200 amino acids, and IFN-λ3 consist of 196 residues. For engineered IFN-λ3 variants, 12 residues were deleted from the wild-type IFN-λ3 sequence, and 12 residues were inserted, resulting in overall length preserved.

As a second example, we analyzed 8 engineered monomeric IL-10 variants. The wild-type human IL-10 consists of 178 amino acids per chain. In the designed group, 11 residues were deleted from the wild-type sequence, 15 residues (for 2 variants), and 20 residues (for 5 variants) were inserted, resulting in a net 4 to 9 residues addition. IL-10M1^44^, a previously developed monomeric IL-10 containing GS linker was also included for comparison.

Finally, the wild-type UGT76G1 and 5 engineered UGT76G1s and their measured T_m_ were collected from previous work^8^. In this group of engineered proteins, 14 to 62 mutations were introduced to the original sequence length of 458 amino acids, including 11 deletions relative to its wild-type. See **Table S5**, **Table S6,** and **Table S7** for measured T_m_ and predicted Δ*G* of IFN-λ, IL-10, and UGT76G1, respectively. The final number of mutations per sequence was recalculated based on the respective sequence alignment with the corresponding wild-type.

Shown in **Fig. 4A**, IFUM prediction results showed a good positive correlation with the experimental T_m_ for all groups of engineered proteins. For IFN-λ, IL-10, and UGT76G1 proteins, PCC of 0.75, 0.62, and 0.87 were observed, respectively, although the results should be interpreted with a caution due to the small sample sizes (p-values: 0.005, 0.104, and 0.023, respectively). For comparison, we selected the AF3 confidence metric (plDDT) (note that all the other ΔΔ*G* prediction methods were not applicable due to length changes). Experimental T_m_ was much less consistent with the AF3 metric, which even negatively correlated with IL-10 and UGT76G1 groups (**Fig. S8**). This highlights the broad generalization of IFUM over AF3 on guiding quite challenging stability engineering problems where both sequence length change and multiple sequence substitution occur simultaneously.

### Practical application 2. Screening *de novo* protein designs for soluble monomeric expression

A common final evaluation step in computational *de novo* protein design is to use a structure prediction model to prioritize candidate sequences^13^. Here, the discriminative power of IFUM, ESMFold, and AF3 are compared for their ability to identify designed proteins that express as soluble monomers in E. coli (**Fig. 4B**). Filtering with IFUM’s Δ*G*_pred_ achieved consistently improved discrimination over the filtering based on plDDT scores from ESMFold and AF3 (**Fig. 4B**, right) across all five different scaffold types: the Rossmann fold^27,28^, TIM barrel^29^, helical repeat^30^, beta barrel^31^, and jelly roll fold^32,33^ (**Fig. 4B**, left, any sequences with sequence identity > 0.30 to the training data were filtered out). This highlights that, similar to its application to protein engineering, IFUM can replace the routinely applied AlphaFold-based metrics in the *de novo* design campaigns to improve design success rates.

## Discussion

### The advantage of explicitly considering unfolded state ensembles

Our results suggest that explicitly considering the unfolded state ensemble contributes to improved accuracy in Δ*G* prediction. This aligns well with the thermodynamic basis of Δ*G* defined as the free energy difference between folded and unfolded states. We have shown this in two ways; first, many ΔΔ*G* predictors relying solely on a single folded state (typically the wild-type structure) revealed their limitations on capturing more complex types of mutations ^16,17^, as supported by our benchmark comparison results (**Fig. 3C)**. Second, the observed performance decrement in IFUM when neglecting the equilibrium ensemble was as significant as that of removing key architectural components seen with ablations (**Table S2**).

### Transfer learning of large pre-trained models can improve model robustness

IFUM predicts Δ*G* and the equilibrium ensemble by integrating sequence and structure embeddings from ProtT5 and ESM-IF1, along with structural input. Leveraging transfer learning from extensive datasets, including 2.1 billion and 45 million protein sequences in BFD and UniRef50 (used to train ProtT5), respectively, and 12 million AlphaFold2-predicted structures (used to train ESM-IF1), likely contributes to mitigating overfitting in IFUM, a strategy also seen in ThermoMPNN^16^. This pre-training approach may offer an advantage over methods like ESM_therm_, which showed significant overfitting when fine-tuning the entire ESM2 model on a single dataset^19^.

### Impact of the input folded state structure quality on IFUM

We showed in **Fig. 2E** that, for natural proteins up to 375 residues, IFUM predictions show much improved correlation with experimental data when the subset of high-confidence AF3 models (plDDT > 90) were used as inputs. A similar trend was observed in every group of engineered proteins, where replacing Rosetta FastRelaxed AF3 models with ESMFold models led to a significant decrease in correlation between measured T_m_ and predicted Δ*G* values (**Fig. S9**). This dependence on input structure quality is likely arising from the nature of the network, which assumes that the input structure representation correctly matches the actual folded state structure. Further analysis highlights the role of the quality of folded state structures (**Fig. S10**). While unrealistically low ΔG values were predicted for wild-type proteins when using poor-quality ESMFold structures (indicated by low pLDDT or pTM scores), substituting these with AF3 structures—more closely resembling the corresponding crystal structures—resulted in more reasonable ΔG predictions. These observations suggest that IFUM prediction is more reliable when supplied with more confident folded structural inputs.

### Potential application for selecting residues to design using per-residue predictions

Because IFUM Δ*G*_pred_ is the sum of per-residue contribution predictions, these residue-wise decomposed values can potentially guide the selection of high-priority residues for stability optimization. To test this idea, sequence redesign on *de novo* designed proteins^47^ using ProteinMPNN^48^ was performed comparing two strategies: 1) full redesign and 2) selective redesign focusing only on residues with negative (i.e. destabilizing) contribution predicted by IFUM. As shown in the **Fig. S11**, selective redesign improved IFUM Δ*G*_pred_ over the original design while full redesign did not. While this result is promising, it might be simply a self-consistent outcome, and hence we expect to experimentally validate this strategy in more depth in our future work.

### Limitations of IFUM

While IFUM demonstrates promising performance in protein stability prediction, several inherent limitations warrant consideration. First, the primary training dataset, Mega-scale, possesses an experimental dynamic range of [–1, 5] kcal/mol, with protein sizes ranging up to 80 residues^20^. This restricted range inherent to the proteolysis-based experimental methodology may constrain IFUM’s capacity for accurate extrapolation beyond these boundaries, and may potentially limit its performance for proteins with exceptional stability or with larger sizes. Second, IFUM’s framework is built on a simplified two-state folding model. The simplified and discrete approximation of ensemble representation in this hypothesis should differ from a more realistic continuous conformational distribution. While this approximation is widely used^20^, it may not fully capture the complexities of the folding landscapes particularly for larger or multi-domain proteins. Third, the network does not apply to membrane proteins and obligate oligomers because it was mainly trained on proteins that are water-soluble and monomeric. We intend to address this in further studies, specifically to predict proteins in their natural conditions. Finally, as IFUM relies on transfer learning from large-scale pre-training, its performance is intrinsically linked to the quality and scope of these foundation models and datasets. While transfer learning likely mitigates overfitting, the model’s standalone utility without this pre-training remains to be fully established. These limitations listed above define key directions for the future model development toward enhancing the scope and accuracy of protein stability predictions.

### Concluding remark: Interpretable predictions and possible applications of IFUM

We have shown IFUM’s strong performance on a broad range of test sets as well as its practical utility in protein *de novo* design and engineering, as demonstrated by the correlation between its Δ*G*_pred_ and experimental T_m_ for engineered proteins with many mutations. Given these results, IFUM can provide guidance to various types of protein design or engineering processes, complementing standard checks like plDDT and PAE score based filtering^13^ (**Fig. 4**, **Fig. S8, Fig. S9**). In summary, IFUM offers a robust method for Δ*G* and ΔΔ*G* prediction and proposes a new concept in the deep-learning architecture design. While these findings are encouraging, further research is needed to fully explore the model’s capabilities and limitations and to guide future development in protein stability engineering and design.

## Methods

### IFUM architecture

IFUM is a transformer-based model designed to predict multiple protein properties, mainly the unfolding free energy (Δ*G*) and the equilibrium ensemble of conformational states. A schematic description of the IFUM model can be found in **Fig. S1**. The model takes three inputs: a sequence embedding of shape [*N*, 1024] generated by ProtT5 embedder, a one-hot encoded, binned pairwise distance map (distogram) of shape [*N*, *N*, 21] representing the folded structure (predicted by ESMFold unless otherwise stated), and a structure embedding of shape [*N*, 512] extracted from ESM-IF1, where *N* is the sequence length of target protein. The IFUM architecture comprises two main components: a main transformer module incorporating triangle multiplicative updates and task-specific simple output multilayer perceptron (MLP).

Initially, IFUM reshapes the sequence embedding and the structure embedding respectively to a common dimension of [*N*, 21] using differential self-attention^34^. These reshaped embeddings are then concatenated into a tensor of shape [2, *N*, 21].

The main module is a modified version of AlphaFold2’s Evoformer, where triangle attention is removed due to its memory demand (but keeping triangle multiplicative updates) and all self-attention layers are replaced with differential self-attention. During the main blocks, the pairwise distogram and the concatenated embeddings are jointly updated through differential self-attention with pair bias and outer product mean as similar to the original Evoformer. IFUM’s main module comprises 11 of these main blocks.

Finally, an MLP with two hidden layers predicts the final output for each objective. Specifically, Δ*G* and sequence recovery are predicted from the main module-updated embeddings, while the equilibrium ensemble is predicted from the main module-updated pairwise distogram. IFUM predicts a protein’s Δ*G* by predicting per-residue Δ*G* contributions and then through an unweighted summation of these contributions. We observed that optimizing destabilizing sequences of a protein based on per-residue contribution results in a higher predicted Δ*G* than naively redesigning the whole sequence, suggesting potential usage of IFUM on *de novo* protein design (**Fig. S11**). IFUM can also predict ΔΔ*G* by subtracting the predicted wild-type Δ*G* from a predicted mutant Δ*G*, obtained from independent runs of the model.

### Equilibrium ensemble modeling

The Flory random coil model is a well-established model for describing the unfolded state ensemble of proteins (Eq. 1).

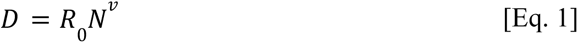

This model provides a statistical description of the average distances expected between residues in a flexible, disordered chain. Therefore, we employed the Flory random coil model with fitted parameters^49,50^ (Eq. 2) to calculate the root-mean-square (RMS) pairwise C𝛼 distances between residue indices *i* and *j* in the unfolded states 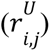 of the protein:

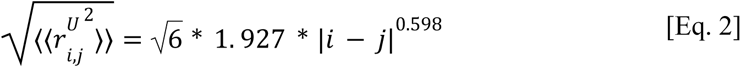

We use these RMS pairwise distances as an explicit representation of the spatial extent of unfolded states to model their contribution to the conformational ensemble.

Protein structure prediction AIs often exhibit insensitivity to structure-disrupting mutations, sometimes predicting structures for mutated sequences that are nearly identical to the wild-type structure. Therefore, we used the ESMFold-predicted structure as a proxy for the target folded (near wild-type) state, recognizing the limitations of AI models in capturing mutational effects. Pairwise C𝛼 distances between residue indices *i* and *j* can be directly calculated from this modeled structure.

Assuming a simple two-state equilibrium between folded and unfolded states, we constructed our model by incorporating the Flory random coil model for unfolded conformations and the ESMFold-predicted structure as a proxy for folded conformations. Within this model, the distance between any two residues was represented as a weighted average of their folded and unfolded states distances. This weighting factor, reflecting the equilibrium ratio between folded and unfolded states (Eq. 3), can be derived directly from the protein’s unfolding free energy, Δ*G* (Eq. 4, note that this is only used at the training stage):

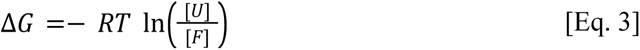

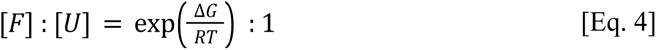

Consequently, using this equilibrium ratio we calculated the ratio of pairwise distances between residues *i* and *j* in folded and unfolded states (Eq. 5):

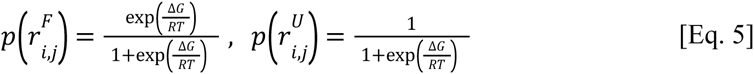

which is then represented as a mixed label distogram (probability values above at *r^F^* and *r^U^* bins otherwise 0) with a range of [2.0, 42.0] Å and a bin width of 2 Å (**Fig. 1B, C**. α corresponds to 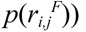. IFUM was trained to predict this equilibrium ensemble distogram.

### Training configuration

IFUM was trained on a single A6000 48 GB GPU using an AdamW with a weight decay of 0.01 and a learning rate of 1 x 10^-4^. The loss function was a weighted combination of three components (Eq. 6, 𝐿):

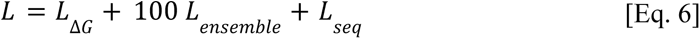

which consist of a Gaussian negative log-likelihood loss for Δ*G* prediction (𝐿_Δ𝐺_), cross-entropy loss for equilibrium ensemble modeling (𝐿_𝑒𝑛𝑠𝑒𝑚𝑏𝑙𝑒_), and a cross-entropy loss for sequence reconstruction (𝐿_𝑠𝑒𝑞_). During training, we used a batch size of 80, increasing it to 100 for validation and testing, and reducing it to 1 for test-only datasets. Data was shuffled at the start of each training epoch, and masks were carefully employed during calculation to enable batch processing of sequences with varying lengths. We acknowledge that we did not explicitly evaluate the impact of different mask sizes on prediction performance.

### Dataset preparation

The Mega-scale dataset was first curated by removing data with deltaG values marked as unreliable (dG_ML = “—”). We then filtered out scrambled sequences with deltaG values greater than 0.5 kcal/mol to avoid potential artifacts caused by aggregation. This curation process resulted in a Mega-scale dataset containing 829,927 sequences, with deltaG values used as the training labels.

The DisProt dataset was curated by collecting sequences shorter than 70 amino acids to enrich for IDPs. Longer sequences in the raw dataset often represented stable proteins with flexible loops, not primarily IDPs. This curation yielded a DisProt dataset with 4,705 sequences and this size of curated subset is noteworthy as it is comparable to, or even larger than, training sets commonly employed in similar studies. Although the DisProt dataset lacks experimental Δ*G* labels, it is a database of proteins known to be disordered or unstable. Therefore, during the training of IFUM, we incorporated the DisProt dataset by modifying the loss function to encourage the model to predict Δ*G* values lower than 0.5 kcal/mol for these sequences. This approach implicitly guides the model to learn features associated with disordered proteins, even in the absence of explicit Δ*G* values for DisProt. We reasoned that by training IFUM to predict low Δ*G* values for sequences known to be disordered, the model would better capture the characteristics of unfolded or unstable protein states.

The CATH database (‘cath-dataset-nonredundant-S20’) was curated by first removing sequences with non canonical amino acids or unknown amino acids or more than two segments (**Fig. S5A**), then collecting sequences shorter than 900 amino acids due to computational limitations when using ESMFold for structure prediction (specifically, CUDA out-of-memory issues encountered). Finally, any CATH sequences with higher sequence identity 0.30 with the Mega-scale test set were filtered out. This curation resulted in a CATH dataset with 6,007 sequences for testing purposes. The SAP values were calculated following the protocol (‘cao_2021_protocol_guide’) from Longxing Cao et al^47^.

The “literature” dataset was first curated by collecting sequences and their corresponding experimental Δ*G* values of wild-type proteins from the S669 dataset’s original 92 papers. The dataset was then filtered to include only entries with experimental pH values between 7.0 and 7.5. Subsequently, we removed homooligomerizing sequences based on UniProt annotations^51^, as well as any sequences with greater than 0.30 sequence identity to those in the Mega-scale train set. This yielded a curated, test-only dataset containing 40 sequences and experimental Δ*G* values.

We also used the same four case studies, p53, myoglobin, staphylococcal nuclease, and T4 lysozyme, as previously employed by ThermoMPNN as test-only datasets.

We collected wild-type UGT76G1 and 7 redesigned sequences and their measured T_m_ values from Seong-Ryeong Go et al^8^. Seong-Ryeong Go et al. used Rosetta based protocol and redesigned the wild-type UGT76G1, enhancing the thermostability and enzymatic activity.

Finally, we collected 413 *de novo* designed sequences from five distinctive folds with their corresponding experimental expression data^27–33^.

### Design workflow of IFN-λ3 and IL-10

To improve structural stability, native loop structures of wild-type IFN-λ3 and IL-10 were computationally redesigned using RFdiffusion^52^, respectively. Among the generated models, those with α-helical linkers were identified using the DSSP algorithm and were chosen for sequence design. For the selected backbones, ProteinMPNN generated amino acid sequences, while keeping the undesigned region fixed. Final sequences of those we measured the T_m_ will be available in our GitHub repository (github.com/HParklab/IFUM). See Jeongwon Yun et al. for more details on protein expression and T_m_ measurement.

### Dataset splitting

We used the MMseqs2 easy-cluster^53^ to cluster the Mega-scale and DisProt datasets with a minimum sequence identity cutoff of 0.30. As each cluster contained a different number of sequences, the clusters were split into train and validation/test sets to ultimately yield an 8:2 ratio of sequences in the final datasets. The validation and test sets were then split randomly in half. The exact values can be found in **Table S8.**

A set of 5,356 sequences, common to both the IFUM Mega-scale test set and the ESM_therm_ test-set-only domains, was assembled and used for IFUM and ESM_therm_ comparison (**Fig. 2D** and **Fig. 3D**; Mega-scale Common).

### Ablation results

All modifications were trained on the same training sets with the same hyperparameters. The best model for each modification was selected based on the highest coefficient of determination (R^2^) on the validation set. All modifications were evaluated with the same test sets to calculate the performance metrics.

### Other predictors

ThermoMPNN is a graph neural network, a transfer-learned version of ProteinMPNN, trained on the Mega-scale dataset to predict ΔΔ*G* from point mutations. It is important to note that ThermoMPNN is designed for point mutations only and does not predict the effects of insertions or deletions, as it requires the wild-type and mutant protein sequences to be of the same length. We generated single mutants ΔΔ*G* values for comparison using the Google Colab implementation of ThermoMPNN. We used ThermoMPNN-D to predict double mutants ΔΔ*G*.

ESM2 is a protein language model trained on the UniRef database to predict masked amino acids in protein sequences. It has demonstrated the ability to predict the functional effects of mutations without requiring any specific training (zero-shot prediction). To quantify this predictive power on predicting Δ*G*, we calculated the ESM2 pseudolikelihood for a sequence of length *N* by summing the log probabilities of predicting each residue 𝑟_i_ when it is masked (Eq. 7):

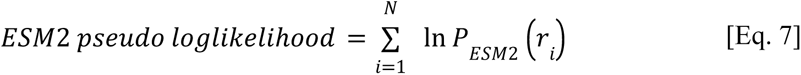

We used esm2_t36_3B_UR50D to generate these pseudo-loglikelihood (plll) values for our comparisons.

ESM_therm_ is a protein language model, a fine-tuned version of ESM2, trained on the Mega-scale dataset to predict Δ*G* of proteins. We used the original script provided in the ESM_therm_ GitHub repository to generate Δ*G* values for our comparisons.

FoldX calculates the Δ*G* value of a protein using energy function. We used predicted structures and the FoldX Stability protocol (FoldX 5.0) to calculate the Δ*G* and ΔΔ*G* values, including indel mutants.

Rosetta uses physical and empirical potentials to calculate the Δ*G* value. We used predicted structures and the FastRelax and Cartesian ddG protocols^39,40^ (ref2015) to calculate the Δ*G* and ΔΔ*G* values, respectively, including indel mutants.

## Acknowledgements

We thank Prof. Junseock Koh, Byung-hyun Bae, Seongin Jung, Soohong Min, and Jinyoung Byun for insightful discussions. This work was supported by the National Research Foundation of Korea (NRF) grant (No. 2022R1C1C1007817 for H.P.), Korea-US Collaborative Research Fund (KUCRF) funded by the Ministry of Science and ICT and Ministry of Health & Welfare (grant number RS-2024-00467483 for H.P.), the Korea Institute of Science and Technology (KIST) Institutional Program (No. 2E33791 for H.P.), and Scale-up TIPS funded by the Ministry of SMEs and Startups (RS-2024-00467110 for H.M.K).

## Contributions

H.L., H.M.K, and H.P. conceived and designed the study. H.L. and H.P. developed the deep learning model. H.L. curated datasets. M.S. reviewed the code. Y.C., J.Y., and H.M.K. conducted protein expression and T_m_ measurement. H.L., H.P., and H.M.K wrote the manuscript. Y.C. and M.S. edited the manuscript. H.L. and M.S. worked on the Google Colab implementation. All authors contributed to the discussion of the results and reviewed the manuscript.

## Additional information

Supplementary information is available.

## Data availability

The code for IFUM is publicly available on GitHub (github.com/HParklab/IFUM) or google colab (https://colab.research.google.com/drive/14TbHFp-BXfiv0vrCSNxyIlqMDOWX-8nV?usp=sharing). All data used in this study can be found in the original research publications: Mega-scale^20^, DisProt^25^, CATH^26^, S669^24^, UBA1^4^, MyUb^5^, Myoglobin^41^, P53^42^, S. nuclease^43^, T4 lysozyme^43^, IFN-λ^14^, UGT76G1^8^, Rossmann fold^27,28^, TIM barrel^29^, helical repeat^30^, beta barrel^31^, and jelly roll fold^32,33^. Every AF3 model structure used for our test sets was generated from the official AlphaFold3 server (www.alphafoldserver.com)^11^.

