## Supplementary Material for "Protein folding stability estimation with explicit consideration of unfolded states"

Table of Contents

1. Supplemental Figures
  - Fig. S1. IFUM main module diagram
  - Fig. S2. Consistency between predicted $\Delta G$ and equilibrium ensemble
  - Fig. S3. Distribution of spatial aggregation propensity (SAP) values in CATH domains grouped by predicted stability
  - Fig. S4. Example of "exposed hydrophobic core" in CATH
  - Fig. S5. Examples of IFUM-predicted $\Delta G < 0$ sequences in CATH
  - Fig. S6. Mega-scale Common: double$_{\Delta\Delta G}$ performance
  - Fig. S7. Relationship between protein stability and melting temperature

2. Supplemental Tables
  - Table S1. Curated 40 unique wild-type proteins from the S669 dataset
  - Table S2. IFUM ablation results on key components
  - Table S3. Characteristics of CATH domains grouped by $\Delta G_{\text{pred}}$
  - Table S4. Mega-scale Common: double$_{\Delta\Delta G}$ detailed performances
  - Table S5. List of experimented IFN-λ
  - Table S6. List of experimented IL-10
  - Table S7. List of experimented UGT76G1
  - Table S8. An overview of collected datasets

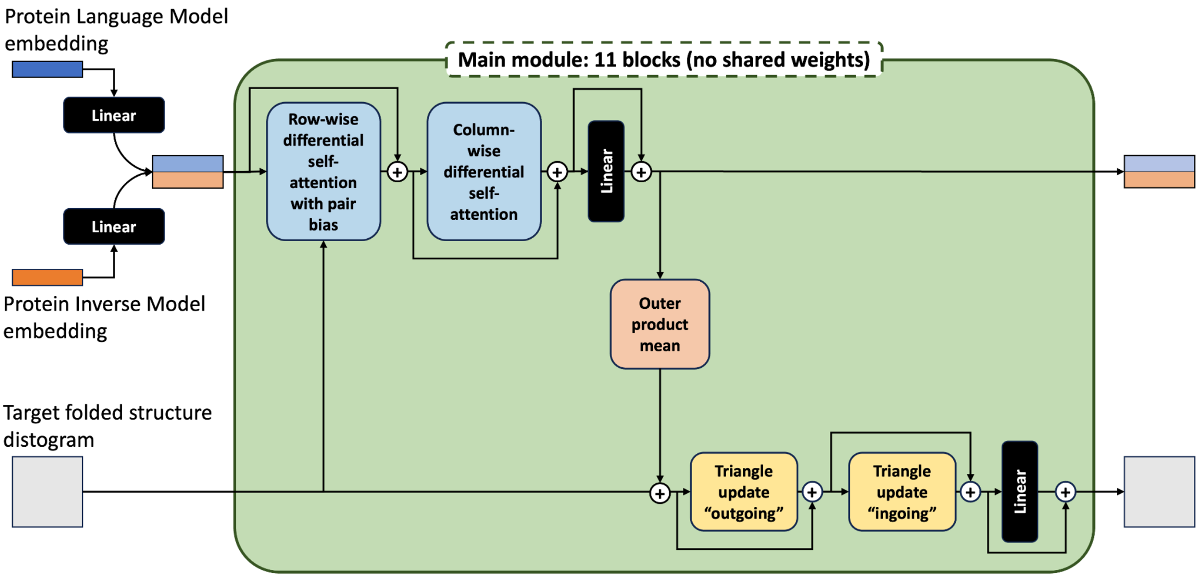


**Fig. S1. IFUM main module diagram.** Diagram that shows how embeddings and distogram are updated in the repetitive main module (green box). The main module is a simplified version of AlphaFold2’s Evoformer[10]. The two major updates were: First, the removal of Evoformer’s original triangle attention layers. Second, replacement of the self-attention layers with differential self-attention. See John Jumper et al. for more details about the Evoformer.

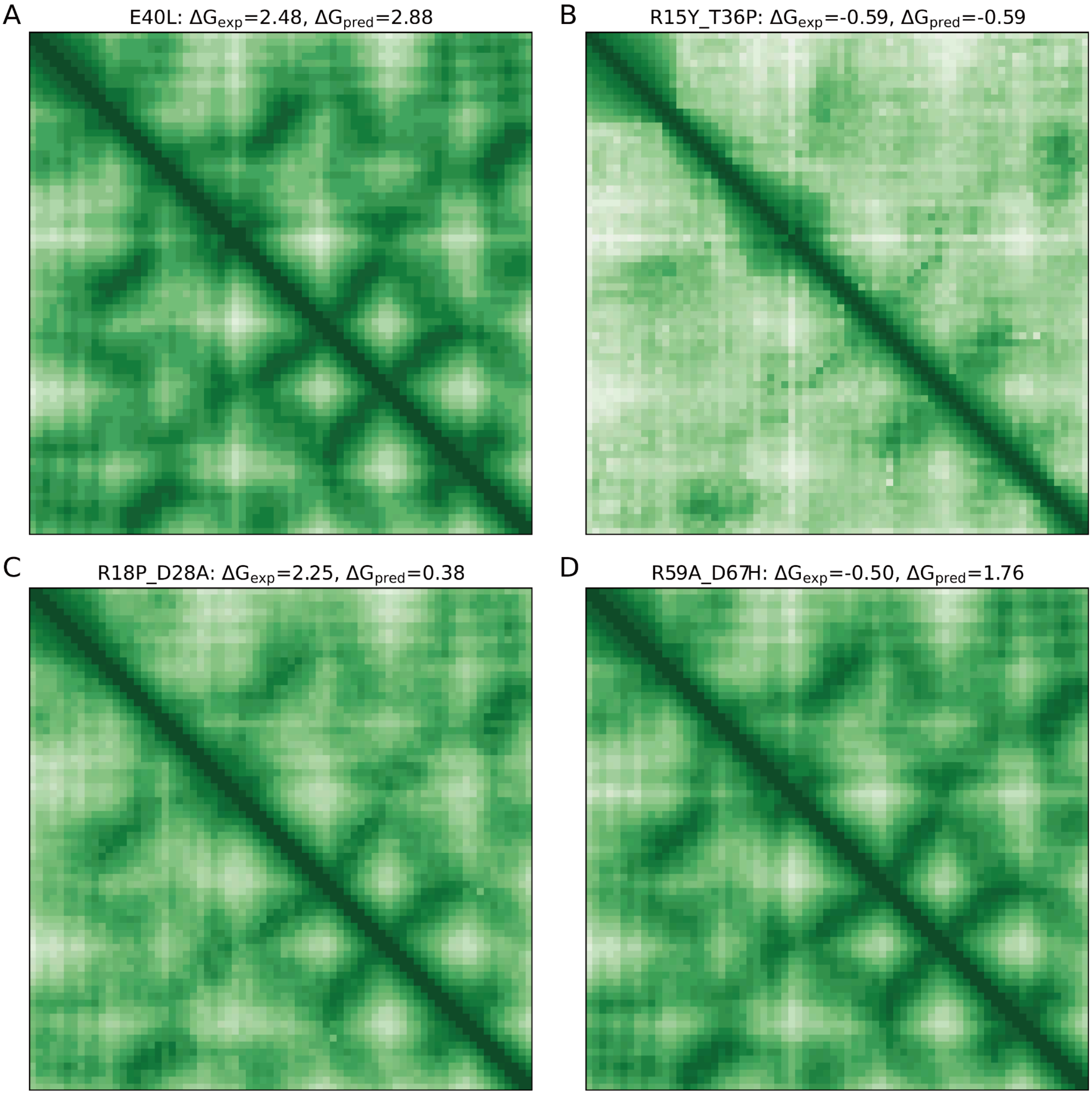


**Fig. S2. Consistency between predicted Δ*G* and equilibrium ensemble.** Four 3DKM mutant examples (E40L, R15Y_T36P, R18P_D28A, R59A_D67H) randomly selected for illustration of the correlation between the predicted Δ*G* and the equilibrium ensemble distograms. The pixel axes indicate the residue indices and the color indicates predicted mean pair distance of the equilibrium ensemble (brighter: higher distance). (A) Represents a case where $\Delta G_{pred}$ is accurate and higher (2.88 kcal/mol vs $\Delta G_{exp}$ = 2.48 kcal/mol). (B) Represents an accurate and lower $\Delta G_{pred}$ (−0.59 kcal/mol vs $\Delta G_{exp}$ = −0.59 kcal/mol). (C) Represents an inaccurately lower $\Delta G_{pred}$ (0.38 kcal/mol vs $\Delta G_{exp}$ = 2.25 kcal/mol). (D) Represents an inaccurately higher $\Delta G_{pred}$ (1.76 kcal/mol vs $\Delta G_{exp}$ = −0.50 kcal/mol).

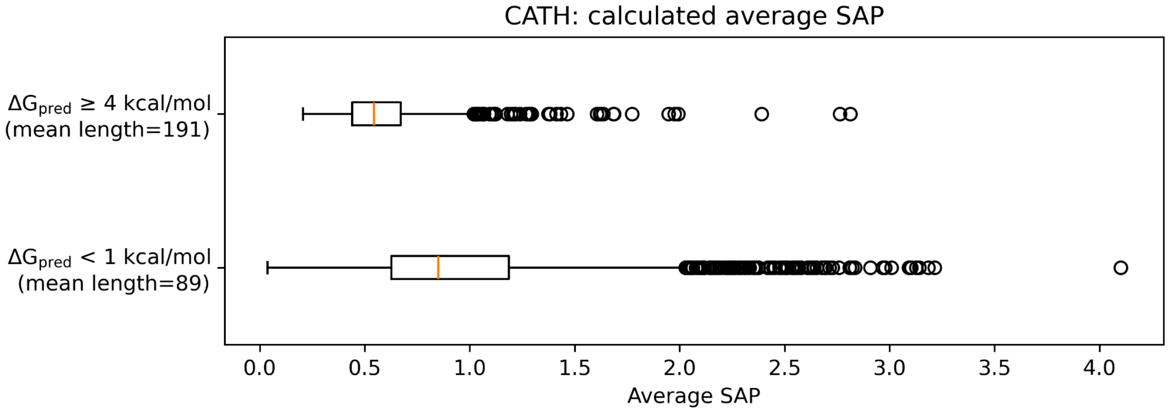


**Fig. S3. Distribution of spatial aggregation propensity (SAP) values in CATH domains grouped by predicted stability.** SAP per amino acid per protein is shown for unstable ($\Delta G_{\text{pred}}$ < 1 kcal/mol) and stable ($\Delta G_{\text{pred}} \geq$ 4 kcal/mol) groups.

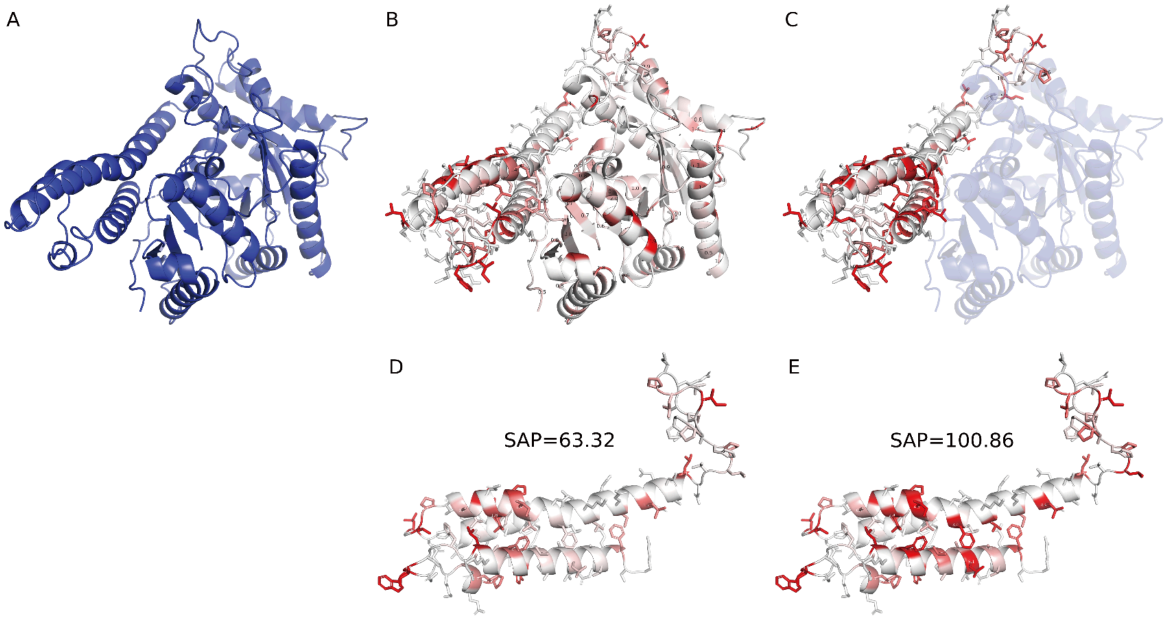


**Fig. S4. Example of "exposed hydrophobic core" in CATH.** (A) A crystal structure of carboxysome shell carbonic anhydrase (PDB: 2FGY). (B-C) Visualization of spatial aggregation propensity (SAP) values for individual amino acids. (B) SAP within the full protein crystal structure of 2FGY (red indicates high SAP, white indicates low SAP). (C) SAP within the CATH domain 2fgyA01. The blue transparent region indicates the portions of the original PDB structure that are absent in this CATH domain. (D) An enlarged view of the 2fgyA01 domain as it appears within the context of the full 2FGY protein structure (as shown in B). Here, the SAP value for this domain within the full protein is indicated as 63.32. (E) An enlarged view of the 2fgyA01 domain when shown in isolation, from the same perspective as (C). This highlights a significant increase in its SAP value (100.86) compared to its SAP value when part of the full protein (63.32 in D). This elevated SAP upon isolation indicates a serious exposure of hydrophobic residues that were previously buried within the intact protein structure.

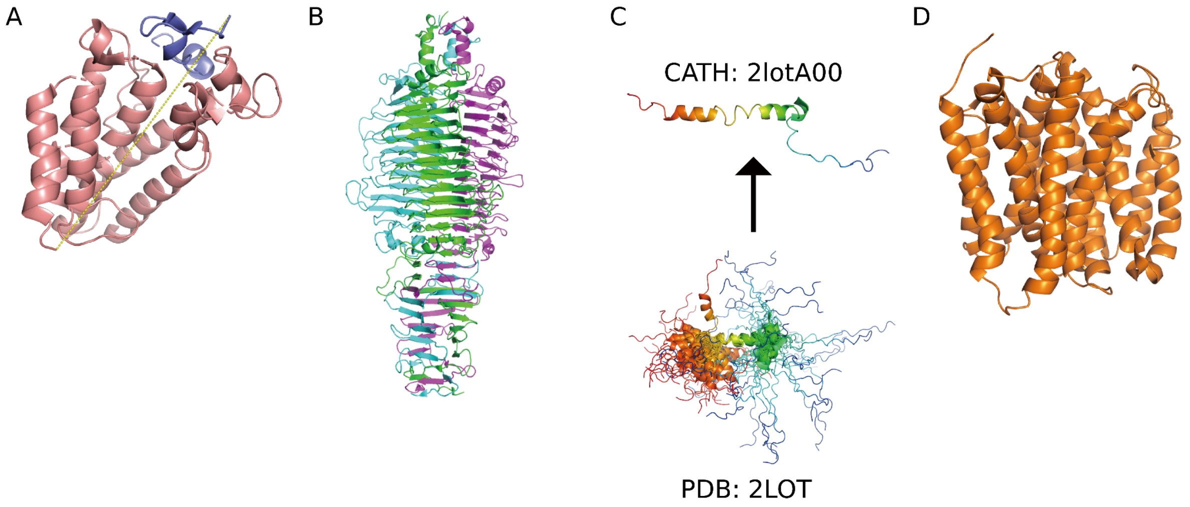


**Fig. S5. Examples of IFUM-predicted Δ*G* < 0 sequences in CATH.** (A) Broken polypeptide bond. CATH domain 3mcxA01 (SusD superfamily protein) comprises two non-contiguous sequence segments (residues 40-254, pink, and 466-492, blue) from a protein. Distance between Cα atoms of residue 254 and 466 is 48.1 Å (yellow dashed line). IFUM inference concatenates these domains in a continuous sequence, forcefully skewing the chain break. Such domains with multiple segments were filtered out and not included in the final 6,007 CATH domains. (B) Obligate oligomer. CATH domain 2x3hA00 (Tail spike protein) represents the monomeric (green) form of an obligate homooligomer protein. The obligate homooligomers were not filtered out. (C) Unfolded protein. CATH domain 2lotA00 (Apelin receptor) is a single state of an unfolded protein, determined by NMR (PDB: 2LOT). NMR structures were not filtered out. (D) Membrane protein. CATH domain 2gfpA00 (Multidrug resistance protein D) is annotated as a membrane protein in UniProt[51], suggesting potential insolubility or aggregation in aqueous solution. Membrane proteins were not filtered out.

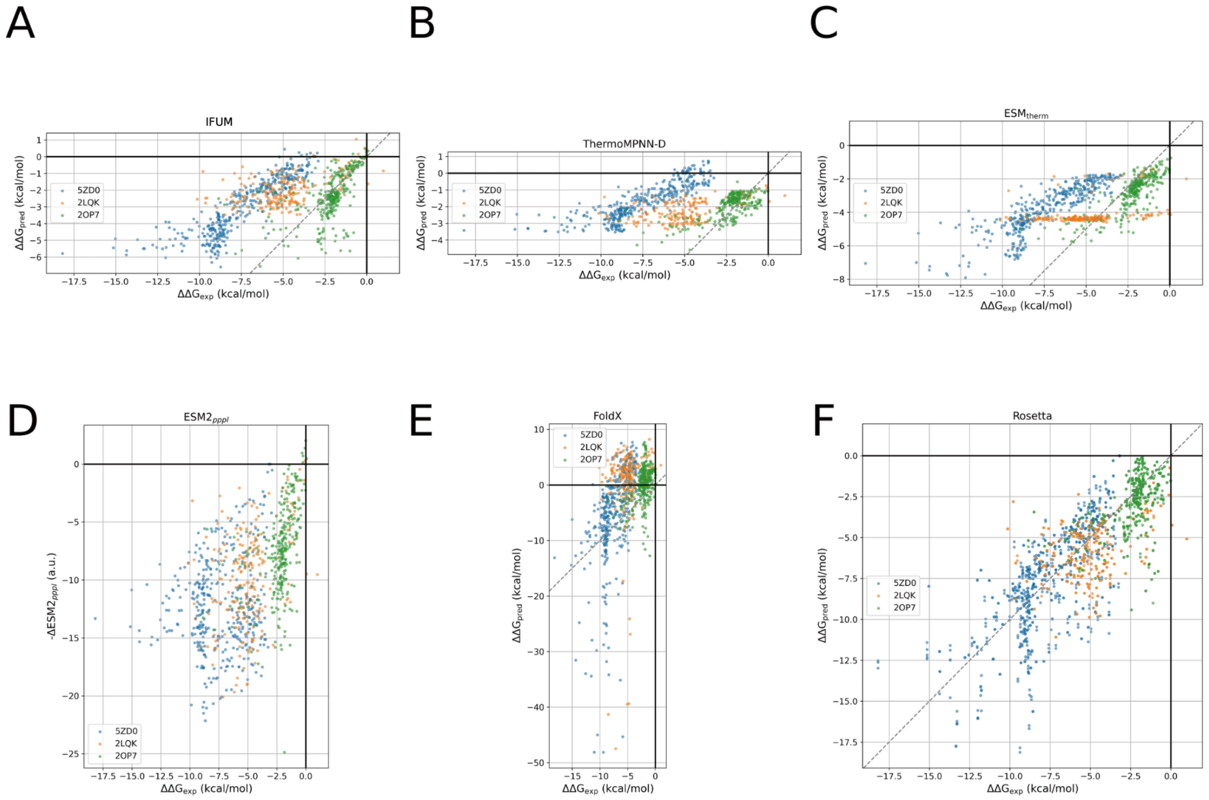


**Fig. S6. Mega-scale Common: double$_{\Delta\Delta G}$ performance.** Scatter plot of each method predicting double mutant $\Delta\Delta G$ values (**Fig. 3C**). (A) IFUM, (B) ThermoMPNN-D, (C) $ESM_{therm}$, (D) $ESM2_{pppl}$, (E) FoldX, and (F) Rosetta (all Rosetta energy values were divided by 2.9, as recommended for the ref2015 score function[39]). The dashed lines indicate perfect correlation (no line for ESM2).

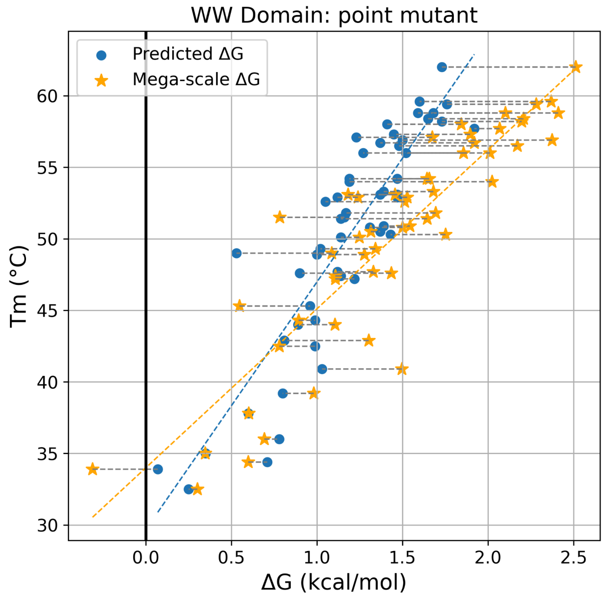


**Fig. S7. Relationship between protein stability and melting temperature.** Scatter plot comparing melting temperature ($T_m$) from Marcus Jäger et al. [54] and $\Delta G$ values are shown (blue: IFUM-predicted, yellow: Mega-scale) for WW domain (PDB: 1I6C). Marcus Jäger et al. performed point mutations on WW domain and measured the $T_m$. While not used for training, the WW domain is part of our validation/test sets. Strong positive correlations were observed between $T_m$ and both predicted and experimental $\Delta G$ values, yielding PCCs of 0.89 and 0.90, respectively. Notably, IFUM predictions and Mega-scale measurements for these WW domain mutants showed a high degree of correlation (PCC = 0.87, RMSE = 0.42 kcal/mol).

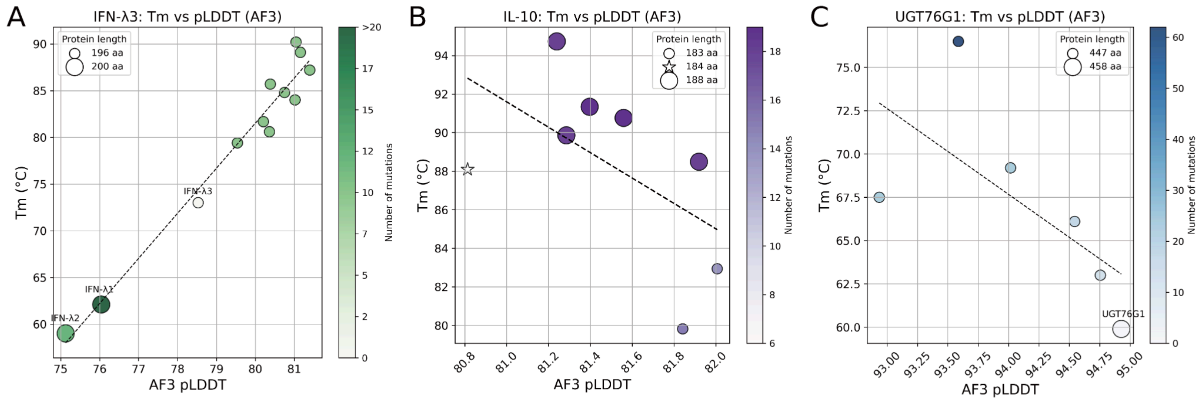


**Fig. S8. Protein engineering: AF3 plDDT and melting temperature ($T_m$).** Scatter plots of $T_m$ versus AF3 plDDT for (A) IFN-λ, (B) IL-10, and (C) UGT76G1. The measured PCC values are 0.98, -0.56, and -0.67, respectively.

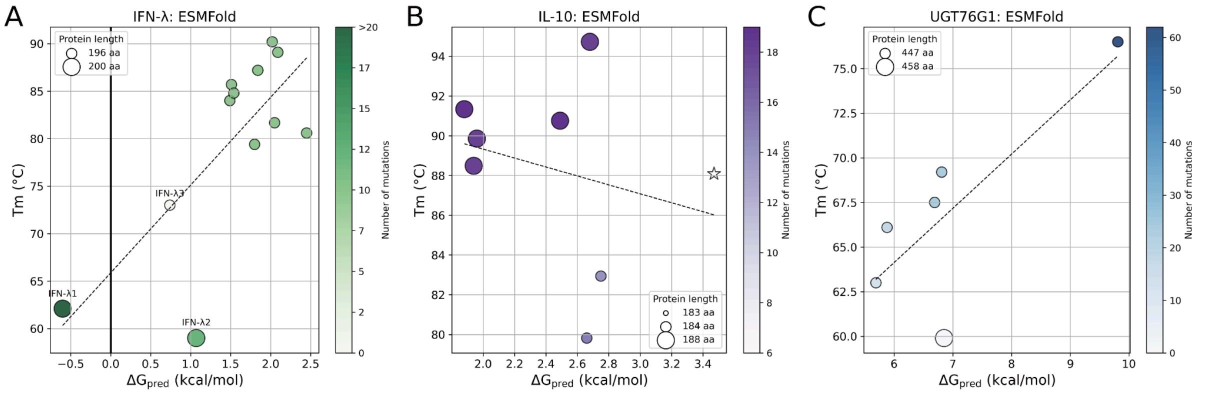


**Fig. S9. Protein engineering: Using ESMFold models as target folded states reduces correlation.** Scatter plots comparing $T_m$ and $\Delta G_{pred}$ are shown for (A) IFN-λ, (B) IL-10, and (C) UGT76G1 using ESMFold models. The Pearson correlation coefficients (PCC) decreased in these plots (IFN-λ: 0.75 to 0.74; IL-10: 0.62 to -0.25; UGT76G1: 0.87 to 0.73) when compared to the results obtained using Rosetta FastRelaxed AlphaFold3 (AF3) models, as presented in **Fig. 4**.

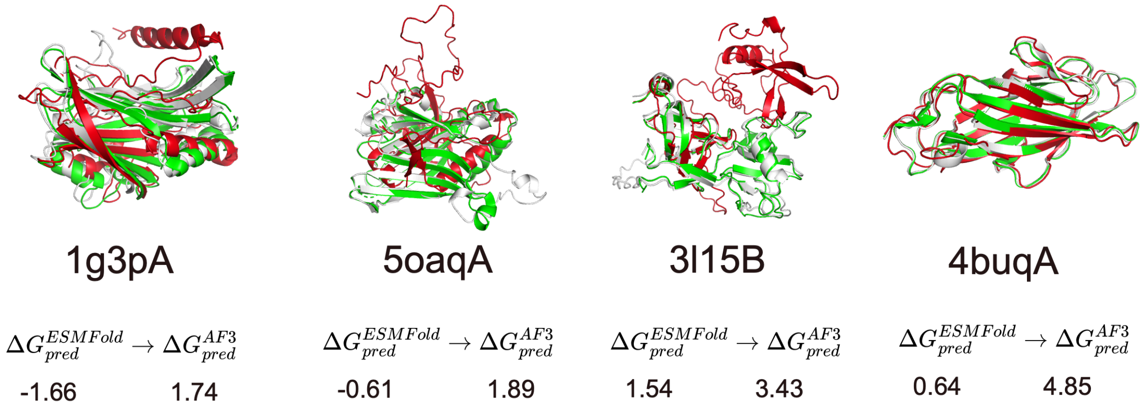


**Fig. S10. Accuracy of IFUM improves when a better target folded state is given.** Wild-type proteins, identified by their PDB IDs, and visualized with crystal structures (green). AlphaFold3 (AF3)-predicted structures (white) are shown to closely resemble these native states. In contrast, ESMFold-predicted structures exhibited instances of poor packing and incorrect topology. In the case of 4buqA, while the overall fold was reasonably captured by ESMFold, loop regions displayed structural inaccuracies. Across all four cases, IFUM's higher $\Delta G$ predictions when provided with improved target folded state inputs (well-packed AF3 structures) suggest effective leveraging of the thermodynamic landscape for interpretable $\Delta G$ estimations. This observation further underscores that IFUM's accuracy in $\Delta G$ prediction is directly linked to the quality of the input representation of the folded state.

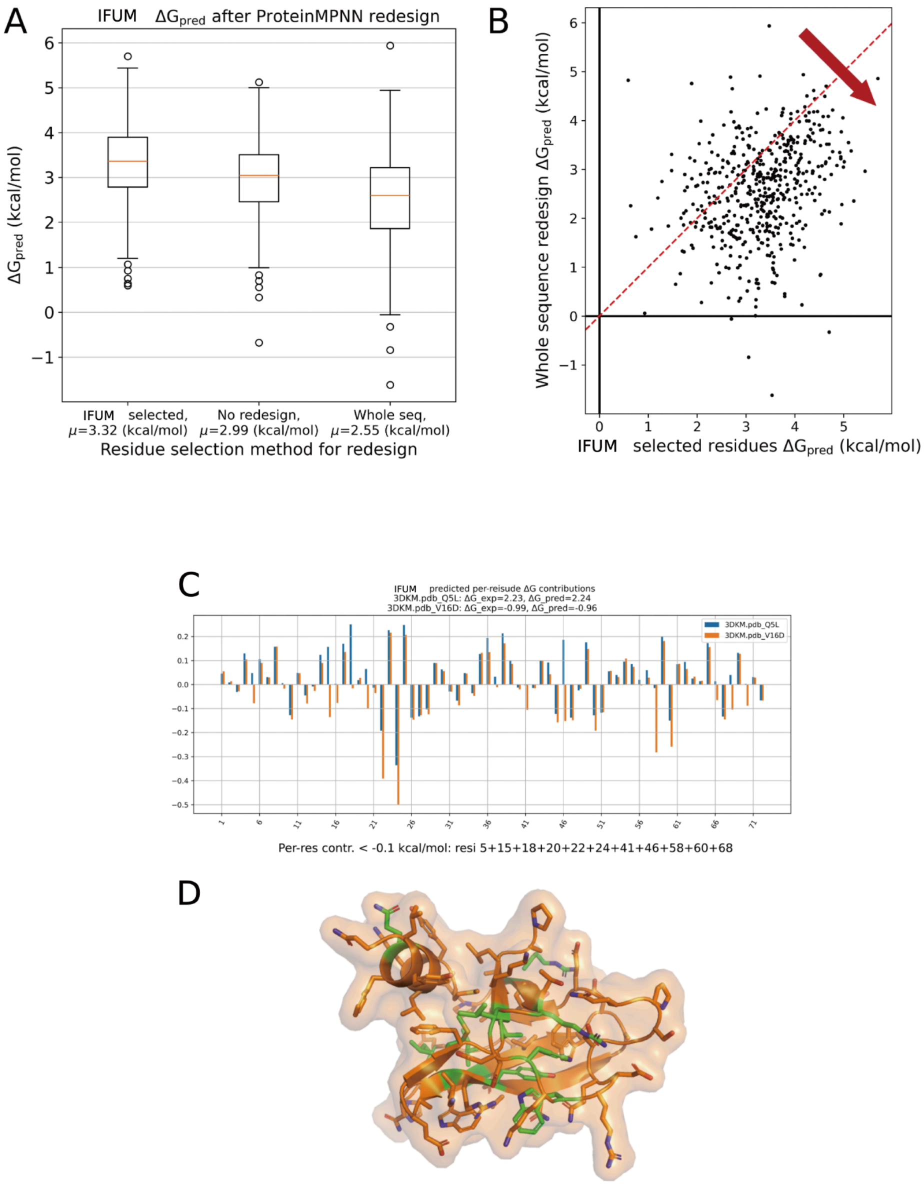


**Fig. S11. Enhancing *de novo* designed protein's $\Delta G_{pred}$ with IFUM guided redesign and per-residue $\Delta G$ visualizations.** The 51 aa long proteins were selected from the Longxing Cao et al. "HHH bc" scaffolds[47], and redesign sequences were generated using ProteinMPNN[48]. (A) Box plots comparing IFUM-predicted $\Delta G$ ($\Delta G_{pred}$) values for 502 *de novo* Rosetta designed three helix bundle proteins using different redesign strategies. Left: IFUM-selected residues. Center: No redesign (original sequences). Right: Whole sequence redesign. (B) Scatter plot comparing the $\Delta G_{pred}$ resulting from whole sequence redesign versus redesign of only IFUM-selected residues from the same starting sequence and structure. Clear shift towards more favorable $\Delta G_{pred}$ values is observed, indicating the effectiveness of IFUM-selected redesign. A red arrow emphasizes this shift. The dashed line indicates equality. (C) Per-residue ΔG Contributions. Bar chart showing per-residue predicted ΔG contributions for wild-type (3DKM.pdb_Q5L) and mutant (3DKM.pdb_V16D) proteins of **Fig. 2C**. (D) Visualization of destabilizing residues. 3D protein structure visualization highlighting residues with per-residue ΔG contributions below –0.1 kcal/mol (green), indicating significant destabilizing residues.

**Table S1. Curated 40 unique wild-type proteins from the S669 dataset.** Name represents the corresponding PDB id and chain id (e.g., 2vy0A represents chain A of a PDB code 2vy0). Sequences and experimental details are available in Corrado Pancotti et al. [24]

| Name | $\Delta G_{exp}$ (kcal/mol) | Name | $\Delta G_{exp}$ (kcal/mol) |
|---|---|---|---|
| 2vy0A | 14.698 | 1fxaA | 6.3 |
| 1xwsA | 12.75 | 1pflA | 6.27 |
| 4buqA | 11.95 | 1bnlA | 5.936 |
| 3d2aA | 11.39 | 1o1uA | 5.9 |
| 1dxxA | 11.2 | 1guaB | 5.721 |
| 3l15B | 9.82 | 3s4mA | 5.6 |
| 5oaqA | 9.7 | 1ft8A | 5.26 |
| 2nteA | 9.6 | 2ltbA | 4.94 |
| 3k82A | 9.321 | 1frdA | 4.5 |
| 1ekgA | 9.1 | 1fh5H | 4.3 |
| 1gwyA | 9.1 | 1itmA | 4.3 |
| 1ir3A | 8.9 | 1xzoA | 4.3 |
| 4bjxA | 8.82 | 4yefA | 4.17 |
| 2ouoA | 8.66 | 1nm1A | 3.94 |
| 4waaA | 8.3 | 1l6hA | 3.895 |
| 2ks4A | 8.21 | 4yeeA | 3.84 |
| 2clrB | 7.887 | 2c9qA | 3.458 |
| 3bciA | 7.189 | 3c2iA | 2.07 |
| 3s92A | 7.14 | 3d3bA | 1.983 |
| 2arfA | 6.453 | 1g3pA | 0.525 |

**Table S2. IFUM ablation results on key components.** Table summarizing performance metrics (RMSE, Pearson correlation coefficient (PCC), Spearman correlation coefficient (SCC), $R^2$) for IFUM and ablated model versions on the same training set, assessing the impact of different model components.

| **Modification** (raw/*clamped*) | **RMSE** (kcal/mol) | **PCC** | **SCC** | $\mathbf{R^2}$ |
|---|---|---|---|---|
| IFUM | **1.59** ***(1.16)*** | **0.78** ***(0.78)*** | **0.74** ***(0.78)*** | **0.54** ***(0.59)*** |
| No equilibrium ensemble ($IFUM_{baseline}$) | 1.73 *(1.39)* | 0.70 *(0.70)* | 0.69 *(0.69)* | 0.45 *(0.40)* |
| No triangle multiplicative update | 1.73 *(1.26)* | 0.69 *(0.69)* | 0.73 *(0.74)* | 0.45 *(0.51)* |
| No differential attention | 1.71 *(1.37)* | 0.71 *(0.71)* | 0.69 *(0.69)* | 0.45 *(0.42)* |

**Table S3. Characteristics of CATH domains grouped by $\Delta G_{pred}$.** Unstable (< 1 kcal/mol) and stable (≥ 4 kcal/mol) CATH domains. For each group, the table reports the total number of domains, the average SAP per residue, and the average protein length. In both average values, Welch's t-test p-values << 0.001.

| $\Delta G_{pred}$ | < 1 kcal/mol | ≥ 4 kcal/mol |
|---|---|---|
| Total number | 2489 | 1659 |
| Average SAP per residue | 1.08 | 0.57 |
| Average protein length | 89 | 191 |

**Table S4. Mega-scale Common: double$_{\Delta\Delta G}$ detailed performances.** The curated double mutant $\Delta\Delta G$ consists of three wild-type domains; 5ZD0, 2LQK, and 2OP7. Pearson Correlation Coefficient (PCC) value per domain, mean value of them, PCC of overall mutants, and the total RMSE are shown. Error canceling effect is reflected as high overall PCC and low mean PCC **(Fig. S6)**.

| **Method** | **5ZD0 PCC** | **2LQK PCC** | **2OP7 PCC** | **Mean PCC** | **Overall PCC** | **Total RMSE** (kcal/mol) |
|---|---|---|---|---|---|---|
| IFUM | **0.87** | **0.51** | 0.55 | 0.65 | 0.61 | 3.51 |
| ThermoMPNN-D | 0.85 | 0.49 | 0.67 | 0.67 | 0.38 | 4.35 |
| $ESM_{therm}$ | 0.86 | 0.39 | **0.77** | **0.68** | 0.70 | 2.79 |
| $ESM2_{pppl}$ | 0.16 | 0.38 | 0.45 | 0.33 | 0.50 | N/A |
| FoldX | 0.55 | 0.11 | 0.40 | 0.35 | 0.50 | 7.12 |
| Rosetta | 0.70 | 0.35 | 0.46 | 0.51 | **0.78** | **2.10** |

**Table S5. List of experimented IFN-λ.** List of IFN-λ proteins with measured melting temperature ($T_m$), IFUM-predicted $\Delta G$ ($\Delta G_{pred}$), and note.

| $T_m$ (°C) | $\Delta G_{pred}$ (kcal/mol) | note |
|---|---|---|
| 80.6 | 1.6 | redesigned IFN-λ3 |
| 79.4 | 1.22 | redesigned IFN-λ3 |
| 81.7 | 1.85 | redesigned IFN-λ3 |
| 90.2 | 1.83 | redesigned IFN-λ3 (DE1) [14] |
| 85.7 | 0.78 | redesigned IFN-λ3 (DE2) [14] |
| 87.2 | 1.35 | redesigned IFN-λ3 (DE3) [14] |
| 84.0 | 1.48 | redesigned IFN-λ3 |
| 89.1 | 2.2 | redesigned IFN-λ3 (DE4) [14] |
| 84.8 | 1.15 | redesigned IFN-λ3 (DE5) [14] |
| 62.1 | -1.0 | wild-type IFN-λ1 [14] |
| 59.0 | 0.70 | wild-type IFN-λ2 [14] |
| 73.0 | 0.93 | wild-type IFN-λ3 [14] |

**Table S6. List of experimented IL-10.** List of IL-10 proteins and their sequence, measured melting temperature ($T_m$), IFUM-predicted $\Delta G$ ($\Delta G_{pred}$), and note.

| $T_m$ (°C) | $\Delta G_{pred}$ (kcal/mol) | note |
|---|---|---|
| 79.8 | 3.22 | redesigned monomeric IL-10 |
| 82.9 | 3.06 | redesigned monomeric IL-10 |
| 94.7 | 3.54 | redesigned monomeric IL-10 |
| 88.5 | 3.75 | redesigned monomeric IL-10 |
| 90.8 | 3.80 | redesigned monomeric IL-10 |
| 89.9 | 3.22 | redesigned monomeric IL-10 |
| 91.3 | 3.72 | redesigned monomeric IL-10 |

**Table S7. List of experimented UGT76G1.** List of UGT76G1 proteins and their sequence, measured melting temperature ($T_m$), IFUM-predicted $\Delta G$ ($\Delta G_{pred}$), and note. Sequences are available in Seong-Ryeong Go et al. [8]

| $T_m$ (°C) | $\Delta G_{pred}$ (kcal/mol) | note |
|---|---|---|
| 59.9 | 6.85 | wild-type UGT76G1 [8] |
| 69.2 | 6.81 | redesigned UGT76G1 (76_4) [8] |
| 76.5 | 9.81 | redesigned UGT76G1 (76_5) [8] |
| 63.0 | 5.69 | redesigned UGT76G1 (76_6) [8] |
| 66.1 | 5.88 | redesigned UGT76G1 (76_7) [8] |
| 67.5 | 6.69 | redesigned UGT76G1 (76_8) [8] |

**Table S8. An overview of collected datasets.**

| Name | Splits | | |
|---|---|---|---|
| | Train | Val | Test |
| Mega-scale | 648,650 | 90,319 | 90,958 |
| DisProt | 3,219 | 694 | 685 |
| CATH | 0 | 0 | 6,007 |
| S669: Wildtypes | 0 | 0 | 40 |
| IFN-λ | 0 | 0 | 12 |
| IL-10 | 0 | 0 | 8 |
| UGT76G1 | 0 | 0 | 6 |
| Protein expression | 0 | 0 | 413 |
